# Diabetes mellitus is associated with increased prevalence of latent tuberculosis infection: Results from the National Health and Nutrition Examination Survey

**DOI:** 10.1101/204461

**Authors:** Marissa M. Barron, Kate M. Shaw, Kai McKeever Bullard, Mohammed K. Ali, Matthew J. Magee

## Abstract

**Aims:** We aimed to determine the association between prediabetes and diabetes with latent TB using National Health and Nutrition Examination Survey data.

**Methods:** We performed a cross-sectional analysis of 2011-2012 National Health and Nutrition Examination Survey data. Participants ≥20 years were eligible. Diabetes was defined by glycated hemoglobin (HbA1c) as no diabetes (≤5.6% [38 mmol/mol]), prediabetes (5.7-6.4% [39-46mmol/mol]), and diabetes (≥6.5% [48 mmol/mol]) combined with self-reported diabetes. Latent TB infection was defined by the QuantiFERON^®^-TB Gold In Tube (QFT-GIT) test. Adjusted odds ratios (aOR) of latent TB infection by diabetes status were calculated using logistic regression and accounted for the stratified probability sample.

**Results:** Diabetes and QFT-GIT measurements were available for 4,958 (89.2%) included participants. Prevalence of diabetes was 11.4% (95%CI 9.8-13.0%) and 22.1% (95%CI 20.5-23.8%) had prediabetes. Prevalence of latent TB infection was 5.9% (95%CI 4.9-7.0%). After adjusting for age, sex, smoking status, history of active TB, and foreign born status, the odds of latent TB infection were greater among adults with diabetes (aOR 1.90, 95%CI 1.15-3.14) compared to those without diabetes. The odds of latent TB in adults with prediabetes (aOR 1.15, 95%CI 0.90-1.47) was similar to those without diabetes.

**Conclusions:** Diabetes is associated with latent TB infection among adults in the United States, even after adjusting for confounding factors. Given diabetes increases the risk of active TB, patients with co-prevalent diabetes and latent TB may be targeted for latent TB treatment.

## 1.1 Introduction

There were 10.4 million incident cases of active tuberculosis (TB) in 2015, and 1.4 million deaths attributable to TB[1]. One-fourth of the global population has prevalent latent tuberculosis infection (LTBI)[2]. Although the lifetime risk of reactivation of LTBI to TB disease only occurs in approximately 10% of infected individuals [3], the risk of progression to TB is higher in individuals with comorbidities, such as HIV[4] and diabetes mellitus[5–11]. Individuals with diabetes have approximately three-times the risk of active TB compared to the general population[6, 8, 10, 12] and 15% of all TB cases are attributed to diabetes[8, 13].

The global diabetes epidemic is steadily increasing[6, 7, 14] and the negative impact of diabetes on TB incidence threatens gains in TB control[15]. In 2014, 415 million adults were living with diabetes, and the prevalence of diabetes is projected to reach 642 million globally by 2040[16]. Additionally, 95% of TB patients reside in low- and middle-income countries, and the largest projected increases in diabetes will occur in these same countries[7, 14].

Although existing evidence has demonstrated a relationship between diabetes and active TB, it is unclear whether diabetes also increases the risk of LTBI. Limited studies that examined the relationship between diabetes and LTBI reported substantial heterogeneity[17] and have not accounted for confounding by clinical comorbidities such as kidney disease or hepatitis[18, 19]. Previous studies that reported an association between diabetes and LTBI were not widely generalizable and mostly have not used reliable measures of diabetes and LTBI[17]. An increased risk of LTBI in patients with diabetes would have major clinical implications for TB and diabetes, especially with the expected increase in global diabetes prevalence. To address the gap in knowledge related to diabetes and LTBI, we aimed to determine the association between prediabetes and diabetes with LTBI using the National Health and Nutrition Examination Survey (NHANES), a study with data that are representative of the US population.

## 1.2 Material and Methods

We conducted a cross-sectional study using NHANES 2011-2012 data, the most recent cycle that includes QuantiFERON^®^-TB Gold In Tube (QFT-GIT) to measure LTBI. Briefly, NHANES is a nationally representative survey of US non-institutionalized civilians that includes an in-person interview followed by a health examination. Details of NHANES methodology have been published previously[20]. In NHANES 2011-2012, 13,431 individuals were selected to participate, 9,756 completed the in-person interview, and 9,338 completed the interview and received an examination[21].

### 1.2.1 Study Design and Participants

Eligible participants were adults (≥20 years) who completed the interview and health examination and had valid QFT-GIT (positive/negative) and diabetes status results. Participants with missing or indeterminate QFT-GIT results were excluded. Participants with missing glycated hemoglobin (HbA1c) and self-reported diabetes status were excluded.

Biological specimen collection was performed in NHANES mobile examination centers (MECs)[20]. Samples were transported to laboratories across the US for processing[22], except samples for QFT-GIT testing, which were processed at a Clinical Laboratory Improvement Act-certified laboratory as previously described[18].

### 1.2.2 Study Measures and Definitions

Diabetes status of participants was defined by self-reported diabetes status and HbA1c. Participants who self-reported previous diabetes diagnosis by a healthcare professional were classified as having diabetes regardless of HbA1c. Participants without self-reported history of diabetes were classified by HbA1c as no diabetes (≤5.6% [38mmol/mol]), prediabetes (5.7-6.4% [39-46mmol/mol]), or diabetes (≥6.5% [48mmol/mol]) according to the American Diabetes Association guidelines[23]. Participants with diabetes were further classified as having diagnosed or undiagnosed diabetes. Diagnosed diabetes was classified as self-reporting diabetes, and undiagnosed diabetes was classified as self-reporting not having diabetes with HbA1c ≥6.5% (48mmol/mol). Among participants with self-reported diabetes, length of time since initial diabetes diagnosis and information regarding diabetes medication were assessed via interview. Self-reported information on use of insulin and oral diabetes agents was collected[24].

QFT-GIT was analyzed according to manufacturer instructions. Results were interpreted according to guidelines from the Centers for Disease Control and Prevention (CDC) for using interferon-gamma release assays (IGRAs)[25]. Participants with positive QFT-GIT results were classified as LTBI positive, participants with negative QFT-GIT results were classified as LTBI negative. Participants with indeterminate QFT-GIT results were classified for this analysis as missing.

Participants who self-reported they had ever been told by a health care professional to have active TB were defined as having a history of active TB. Body mass index (BMI) ranges were categorized as underweight (<18.5), normal weight (18.5-24.9 kg/m^2^), overweight (25.0-29.9 kg/m^2^), or obese (≥30 kg/m^2^) according to CDC guidelines[26]. Age ranges were categorized as young adult (20-34 years), middle-aged (35-64 years), or elderly (65 years and older). Current smokers were defined as participants who self-reported use of 100 cigarettes in their lifetime and self-reported currently smoking, which included smokers reporting use every day or some days. Former smokers were defined as those who reported smoking 100 cigarettes in their lifetime but did not currently smoke cigarettes. Participants who had not smoked 100 cigarettes in their lifetime were defined as never smokers [24, 27].

Urine samples were analyzed for albumin creatinine ratio (ACR), and ACR levels were categorized as normal to mildly increased (<30mg/g), moderately increased (30-300mg/g), or severely increased (>300mg/g) according to National Kidney Foundation guidelines for albuminuria categories in chronic kidney disease[28]. Hepatitis B virus (HBV) core antibody (anti-HBc) and surface antigen (HBsAg) response were determined using the VITROS Anti-HBc assay and HBsAg assay, respectively; results were defined as positive or negative. The HBsAg assay was only performed for participants that tested positive for anti-HBc; participants with a negative result for anti-HBc were defined as negative for HBsAg. Hepatitis C antibody (anti-HCV) response was determined using the VITROS Anti-HCV assay; results were defined as positive or negative. Non-fasting blood samples were analyzed for total cholesterol and high-density lipoprotein (HDL) cholesterol[24]. Total cholesterol was defined as desirable (<200mg/dL), borderline high (200-239mg/dL), or high (≥240mg/dL) according to National Institutes of Health guidelines[29]. Our study categorized HDL cholesterol levels as major risk factor for heart disease (<40mg/dL), borderline (40-59mg/dL), or protective against heart disease (≥60mg/dL) according to Medline Plus guidelines[30]. Responses of “don’t know” or “refused” were recoded as missing for all variables.

### 1.2.3 Statistical Analyses

To examine the association between diabetes and LTBI we used bivariate analyses and multivariable logistic regression. The Rao-Scott chi-square test was used to analyze all bivariate associations between participant characteristics and LTBI and diabetes. To examine the prevalence of diabetes and LTBI in the US population, we reported weighted prevalence estimates and 95% confidence intervals (CI). Taylor series method was used to estimate variance for all prevalence estimates[31]. Multivariable logistic regression was used to estimate adjusted odds ratios (aOR) and 95% CI between diabetes and LTBI and were adjusted for potential confounders. Covariates included in multivariable models as confounders were chosen from observed bivariate associations with diabetes and LTBI, previous study findings, and causal model theory (directed acyclic graphs) [32]. In multivariable models, multiplicative statistical interaction was assessed to determine if the association between diabetes and LTBI was modified by obesity or HDL cholesterol. In a subgroup analysis, we also examined the bivariate association between participant characteristics and LTBI only among individuals with diabetes. Analyses were performed using SAS version 9.4 and accounted for the weighted stratified probability sample design of NHANES [33]. Because medical examination data were used during the analyses, we used the weight variable WTMEC2YR to obtain prevalence estimates and measures of association. A two-sided p-value <0.05 was considered statistically significant for all tests.

### 1.2.4 Sensitivity Analysis

We performed sensitivity analyses to assess potential error due to 1) misclassification of diabetes status and 2) covariate misspecification in multivariable models. To assess diabetes misclassification, we re-examined the diabetes-LTBI association after adding fasting blood glucose (prediabetes 100-125mg/dL, or diabetes ≥126mg/dL) to our primary diabetes definition which used self-report and HbA1c only[23]. In the second sensitivity analysis we specified several subsets of adjusted multivariable models to provide a range of plausible aORs and 95%CI for the association between diabetes and LTBI.

## 1.3 Results

### 1.3.1 Study Participants

Of 9,756 NHANES 2011-2012 participants, 5,560 (57.0%) were aged 20 years or older and thus eligible for our study. A total of 4,958 (89.2%) participants had both valid QFT-GIT results and information on self-reported diabetes status and/or HbA1c results and were included in these analyses. Before accounting for selection weights, 793 eligible participants had diabetes, 513 had LTBI, and 127 had both diabetes and LTBI.

### 1.3.2 Prevalence of Diabetes and Latent TB Infection

Our results estimated that the prevalence of diabetes among adults in the United States population was 11.4% (95%CI 9.8-13.0%) and the prevalence of prediabetes was 22.1% (95%CI 20.5-23.8%) (Table 1). The prevalence of diabetes was highest among the elderly (22.9%; 95%CI 19.8-25.9%), people with obesity (20.4%; 95%CI 17.3-23.5%), those with less than a 9^th^ grade education (25.2%; 95%CI 18.2-32.2%), severely increased ACRs (46.9%; 95%CI 33.7-60.1%), hepatitis C (19.8%; 95%CI 7.1-32.5%), and hypertension (23.3%; 95%CI 20.8-25.9%).

**Table 1:**
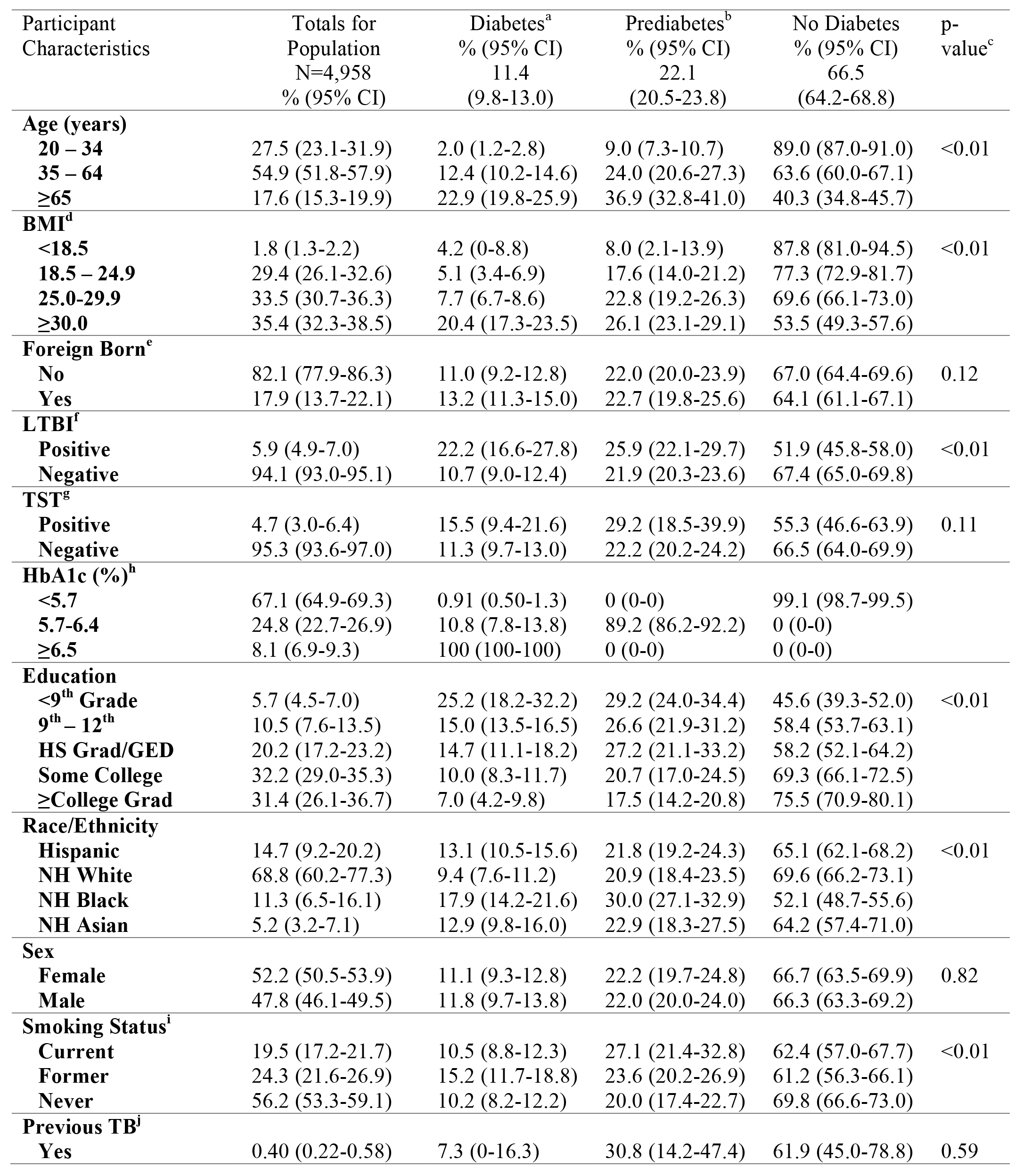
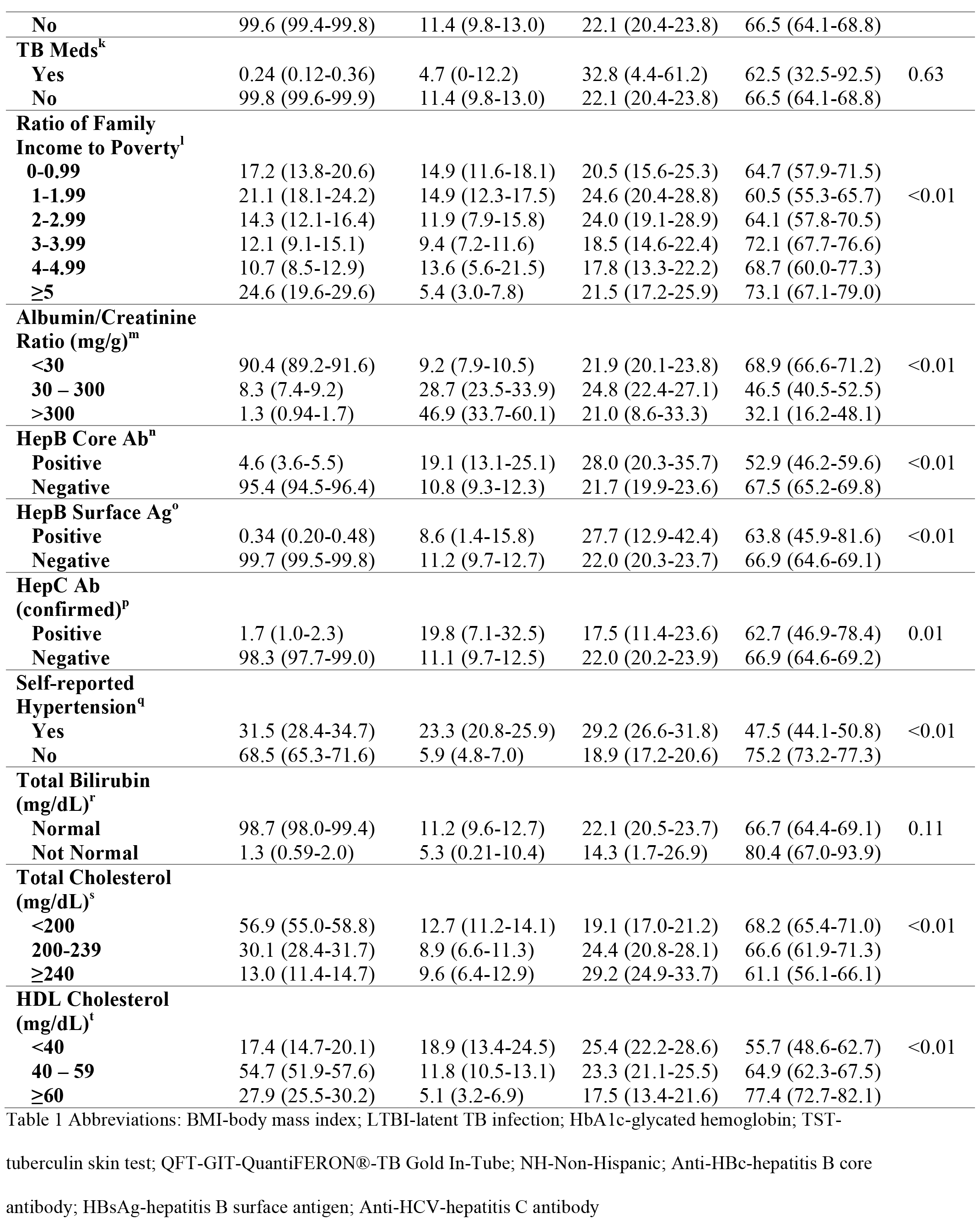
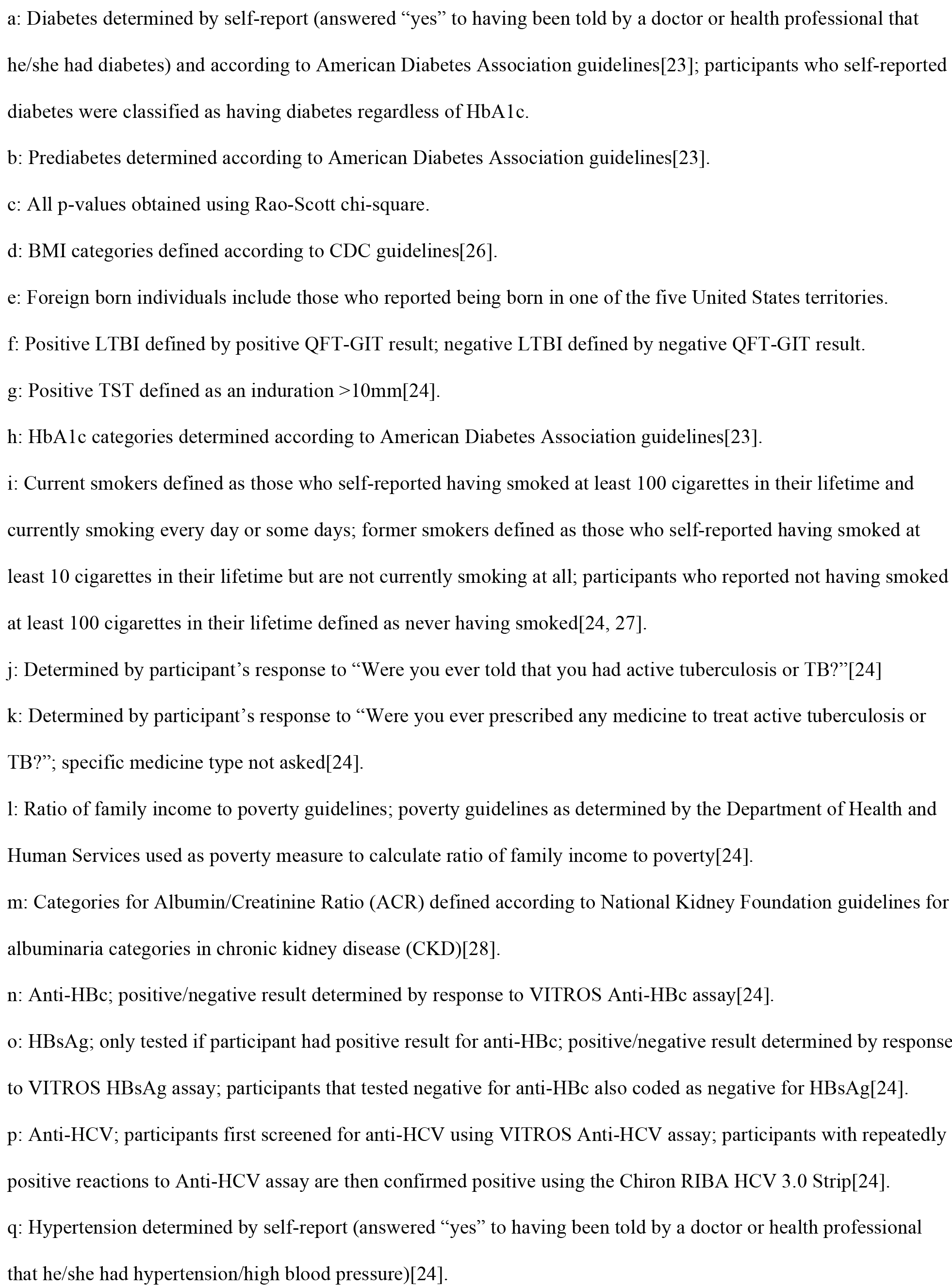
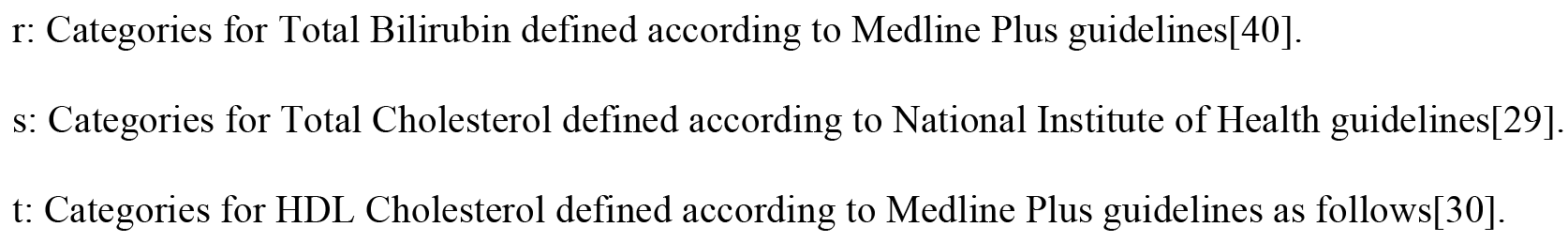
Weighted prevalence of diabetes and prediabetes among the civilian, non-institutionalized United States population, adults 20 years and older, 2011-2012

Our results estimated that the prevalence of LTBI among adults in the United States was 5.9% (95%CI 4.9-7.0%) (Table 2). Prevalence of LTBI was highest among the elderly (8.8%; 95%CI 6.6-10.9%), the foreign-born (17.2%; 95%CI 14.3-20.0%), those with less than a 9^th^ grade education (17.9%; 95%CI 13.2-22.7%), Hispanics (12.9%; 95%CI 10.4-15.4%), non-Hispanic Asians (20.3%; 95%CI 16.8-23.8%), and those who reported a previous history of active TB (42.7%; 95%CI 24.1-61.2). LTBI prevalence was also high among those with high (>300mg/g) ACR (12.9%; 95%CI 6.7-19.2%), those who tested positive for anti-HBc (18.0%; 95%CI 11.6-24.4%), and those who tested positive for HBsAg (23.0%; 95%CI 8.2-37.7%).

**Table 2:**
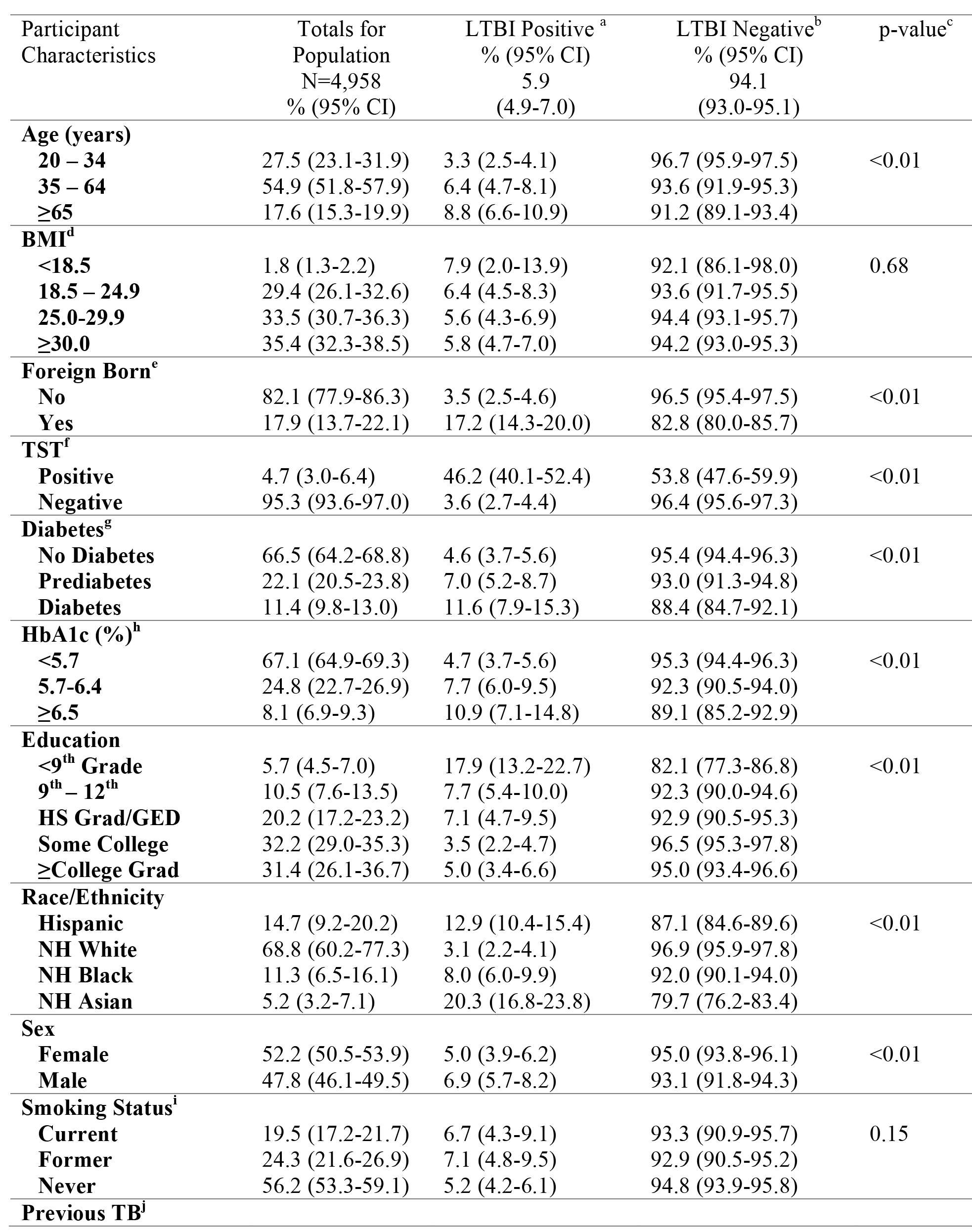
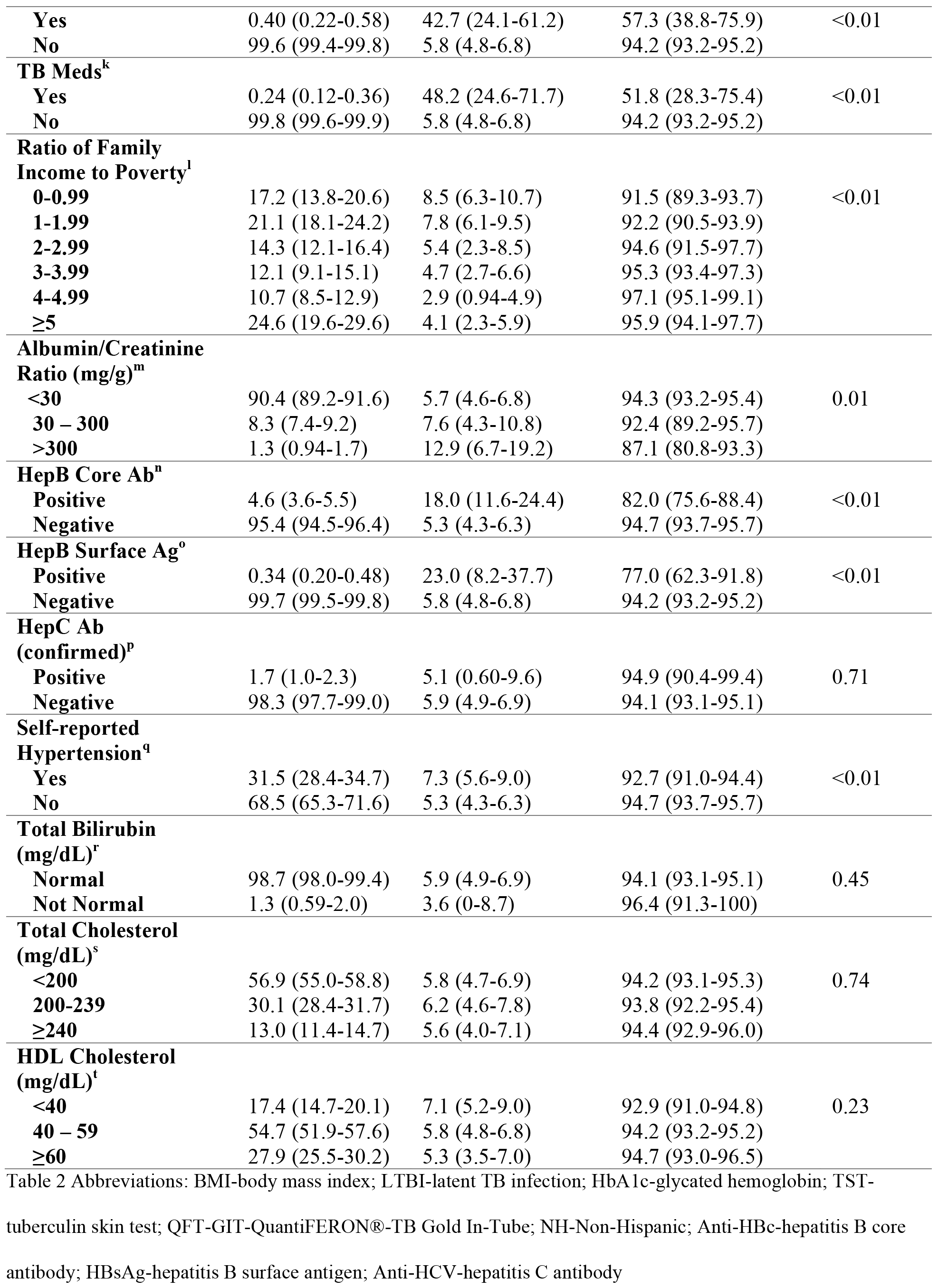
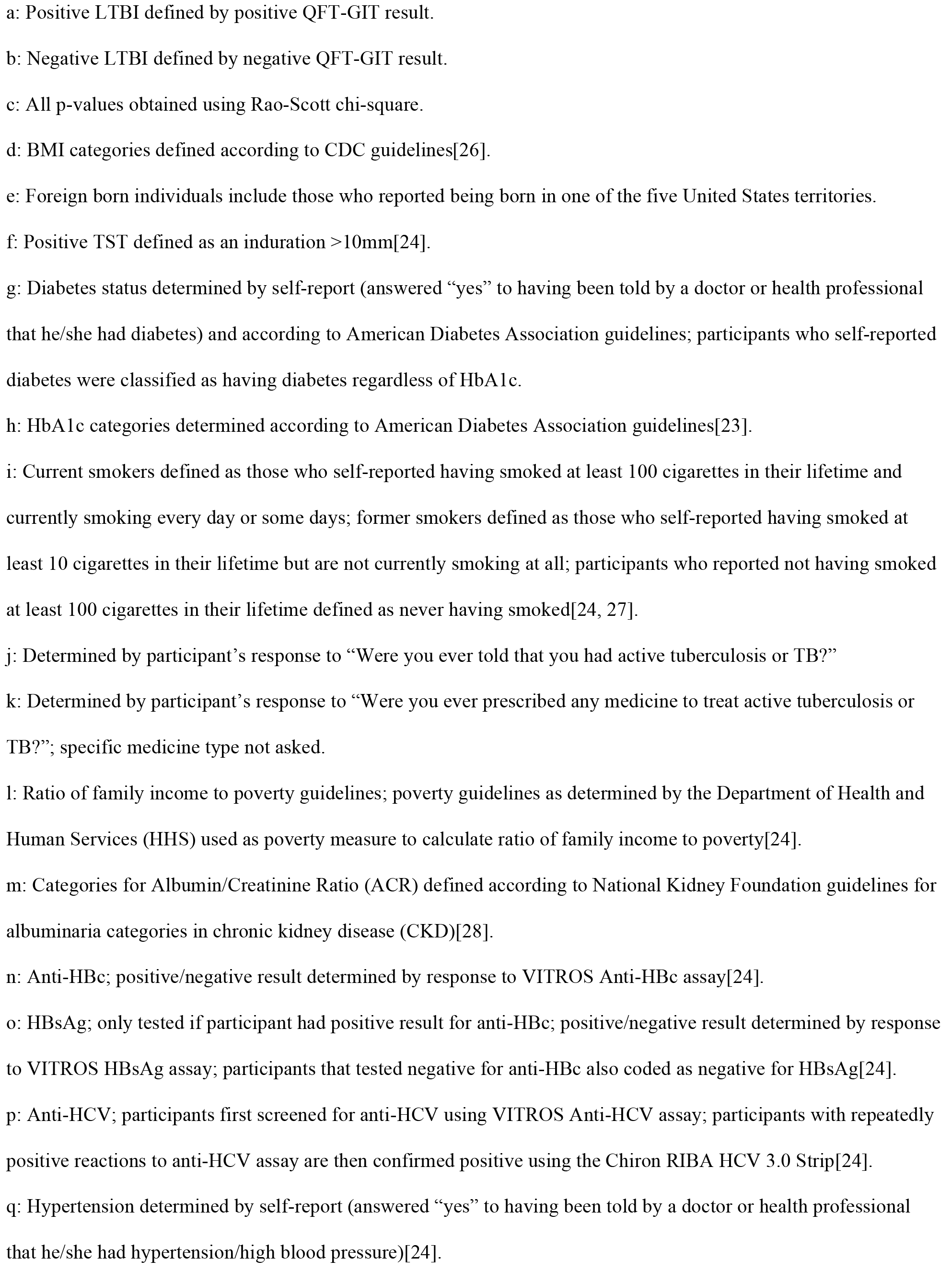
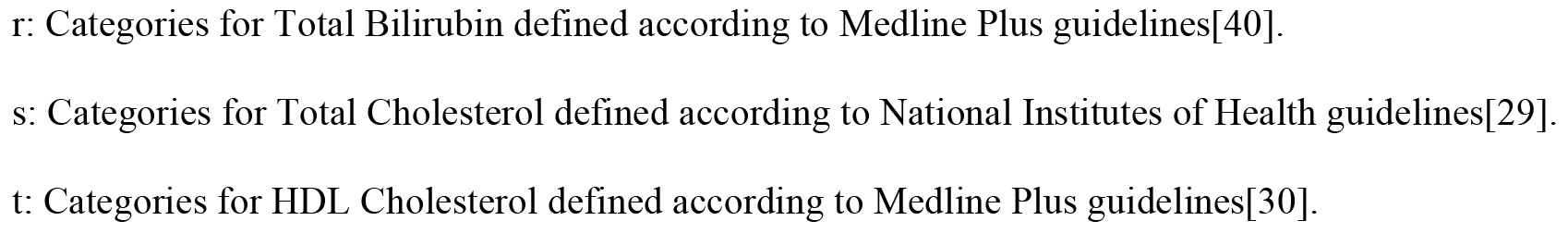
Weighted prevalence of latent tuberculosis infection (LTBI) among the civilian, noninstitutionalized United States population, adults 20 years and older, 2011-2012

The prevalence of LTBI was significantly higher among adults with diabetes (11.6%; 95%CI 7.9-15.3%) compared to those without diabetes (4.6%; 95%CI 3.7-5.6%). LTBI prevalence was also higher among those with prediabetes (7.0%; 95%CI 5.2-8.7%) compared to those without diabetes, though the difference was not statistically significant. Adults with diabetes and prediabetes had significantly higher crude odds of LTBI (diabetes: crude OR 2.70; 95%CI 1.76-4.14; prediabetes: crude OR 1.54; 95%CI 1.24-1.91) compared to those without diabetes (Table 3). Reported inversely, among those with LTBI the prevalence of diabetes was 22.2% (95%CI 16.6-27.8%) and the prevalence of prediabetes was 25.9% (95%CI 22.1-29.7%). Among those without LTBI the prevalence of diabetes was 10.7% (95%CI 9.0-12.4%) and the prevalence of prediabetes was 21.9% (95%CI 20.3-23.6%).

**Table 3:** Multivariable models for odds of latent tuberculosis infection based on positive QFT-GIT result by diabetes status in the civilian, non-institutionalized United States population aged 20 years and older, 2011-2012

| Models | Crude Odds Ratio (95% CI) | Adjusted Odds Ratio (95% CI)<sup>a</sup> |
| --- | --- | --- |
| Model 1: |  |  |
| No Diabetes<sup>b</sup> | 1.00 | 1.00 |
| Prediabetes | 1.54 (1.24 – 1.91) | 1.15 (0.90 – 1.47) |
| Diabetes | 2.70 (1.76 – 4.14) | 1.90 (1.15 – 3.14) |
| Model 2: |  |  |
| No Diabetes<sup>c</sup> | 1.00 | 1.00 |
| Diabetes | 2.38 (1.58 – 3.59) | 1.80 (1.14 – 2.83) |
| Model 3: |  |  |
| No Diabetes | 1.00 | 1.00 |
| Undiagnosed Diabetes<sup>d</sup> | 3.06 (1.61 – 5.82) | 1.96 (1.05 – 3.68) |
| Diagnosed Diabetes | 2.22 (1.46 – 3.38) | 1.75 (1.09 – 2.80) |
| Model 4: HbA1c (%)<sup>e</sup> |  |  |
| <5.7% | 1.00 | 1.00 |
| 5.7-6.4% | 1.71 (1.33 – 2.19) | 1.30 (0.96 – 1.75) |
| ≥6.5% | 2.50 (1.64 – 3.81) | 1.67 (1.04 – 2.69) |
| Model 5: HbA1c (%) |  |  |
| <5.7 | 1.00 | 1.00 |
| 5.7-6.4 | 1.71 (1.33 – 2.19) | 1.30 (0.96 – 1.75) |
| 6.5-7.5 | 2.44 (1.48 – 4.01) | 1.66 (0.99 – 2.80) |
| 7.6-8.5 | 3.12 (1.62 – 6.02) | 1.96 (1.06 – 3.63) |
| >8.5 | 2.21 (1.08 – 4.51) | 1.50 (0.70 – 3.22) |

a: Models adjusted for age (categorized as 20-35 years, 35-65 years, and ≥65 years), sex (female, male), smoking status (current, former, never), history of active TB (yes, no), and foreign born status (yes, no).
b: Diabetes status determined by self-report (answered “yes” to having been told by a doctor or health professional that he/she had diabetes) and according to American Diabetes Association guidelines[23]; participants who self-reported diabetes were classified as having diabetes regardless of HbA1c.
c: Individuals classified as having prediabetes or no diabetes for Model 1 were classified as not having diabetes for Model 2.
d: Participants with diabetes who were unaware they had diabetes were classified as previously undiagnosed; these previously undiagnosed participants have no information for duration of diabetes[24].
e: Glycated hemoglobin (HbA1c) categories determined according to American Diabetes Association guidelines[23].
**Bold** indicates that the adjusted odds ratio (aOR) is statistically significant

### 1.3.3 Multivariable Models Results

Adults with diabetes had significantly higher odds of LTBI (aOR 1.90; 95%CI 1.15-3.14) compared to adults without diabetes (Table 3) after adjusting for age, sex, smoking status, history of active TB, and foreign born status. Those previously diagnosed with diabetes had significantly higher odds of LTBI (aOR 1.75; 95%CI 1.09-2.80) compared to adults without diabetes, as did adults with previously undiagnosed diabetes (aOR 1.96; 95%CI 1.05-3.68). The odds of LTBI among adults with prediabetes (aOR 1.15; 95%CI 0.90-1.47) was not significantly higher than among adults without diabetes.

We found no indication of significant multiplicative interaction. Although not significantly different from each other, the association between diabetes and LTBI tended to be greater among those with obesity (aOR 2.22; 95%CI 1.08-4.54) compared to those without obesity (aOR 1.48; 95%CI 0.85-2.58) (data not shown). Similarly, the association between diabetes and LTBI was non-significantly greater among those with higher HDL (≥60mg/dL) levels (aOR 2.77; 95%CI 1.13-6.84) compared to those with lower HDL (<60mg/dL) levels (aOR 1.80; 95%CI 0.99-3.29).

### 1.3.4 Subgroup Analysis of Adults with Diabetes

Among adults with diabetes, an estimated 19.9% (95%CI 15.3-24.4%) were previously undiagnosed (Table 4). Prevalence of LTBI was non-significantly (p-value=0.24) different among adults with previously undiagnosed diabetes (14.4%; 95%CI 6.7-22.2%) compared to adults who had been previously diagnosed (10.9%; 95%CI 7.4-14.4%). LTBI prevalence was significantly higher (p-value=0.03) among adults who reported not using insulin (12.9%; 95%CI 8.5-17.3%) compared to adults who reported using insulin (7.3%; 95%CI 3.1-11.4). Among those with diabetes, LTBI prevalence was found to be highest among Hispanics (24.3%; 95%CI 12.4-36.2%), non-Hispanic Asians (27.5%; 95%CI 19.0-35.9%), those born outside of the United States (30.2; 95%CI 18.8-41.6%), and those with a positive test result for anti-HBc (20.8%; 95%CI 8.9-32.7%).

**Table 4:**
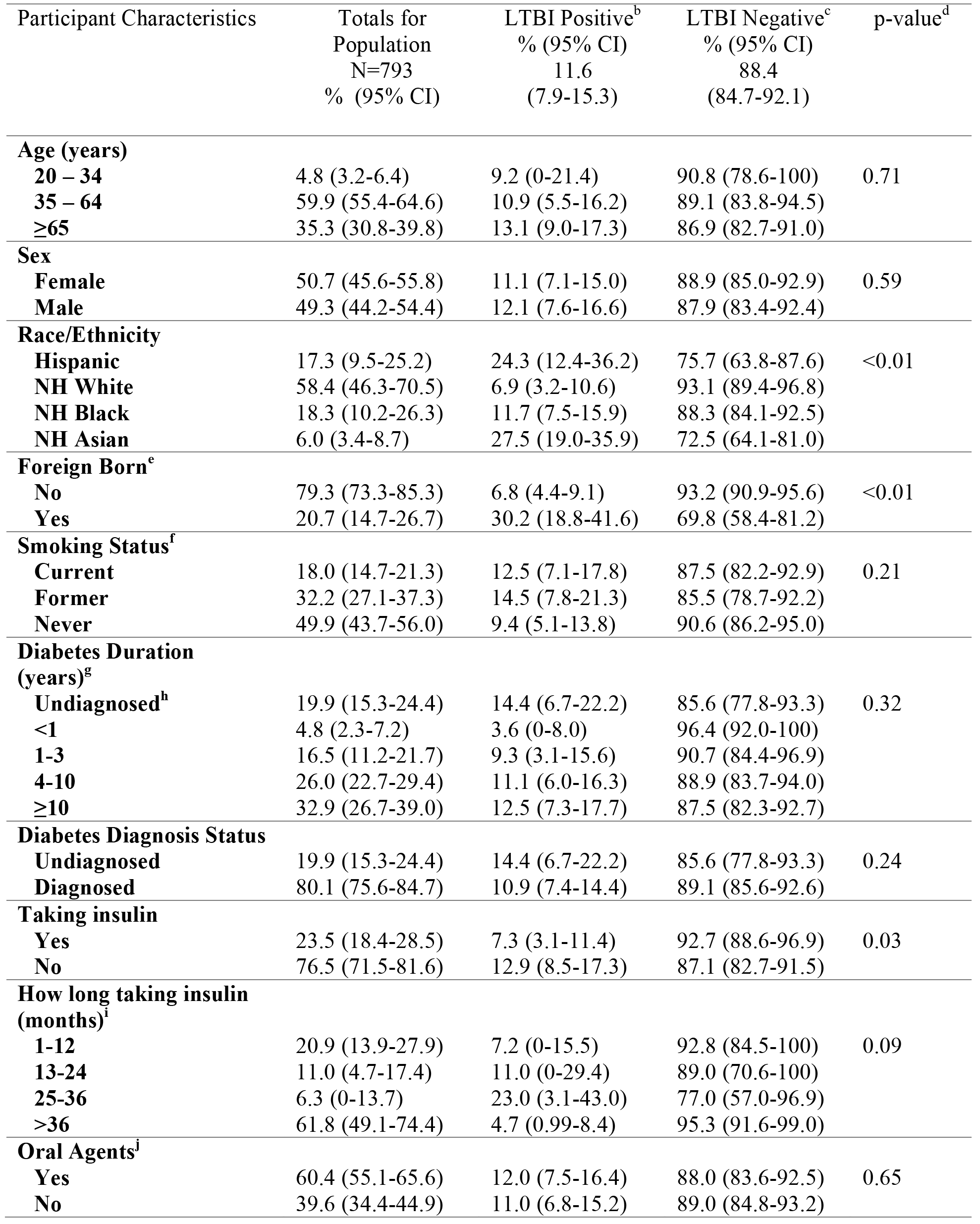
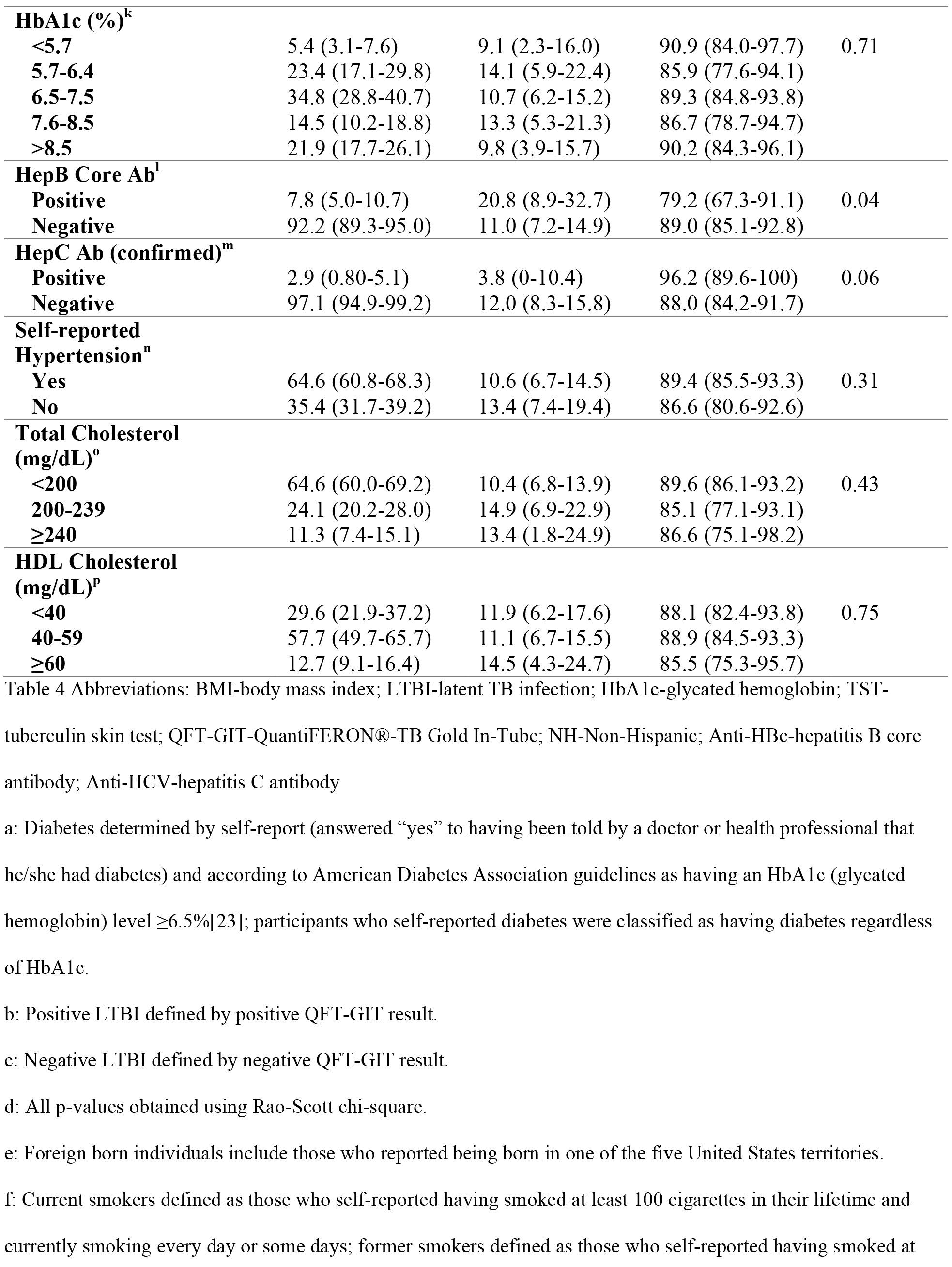
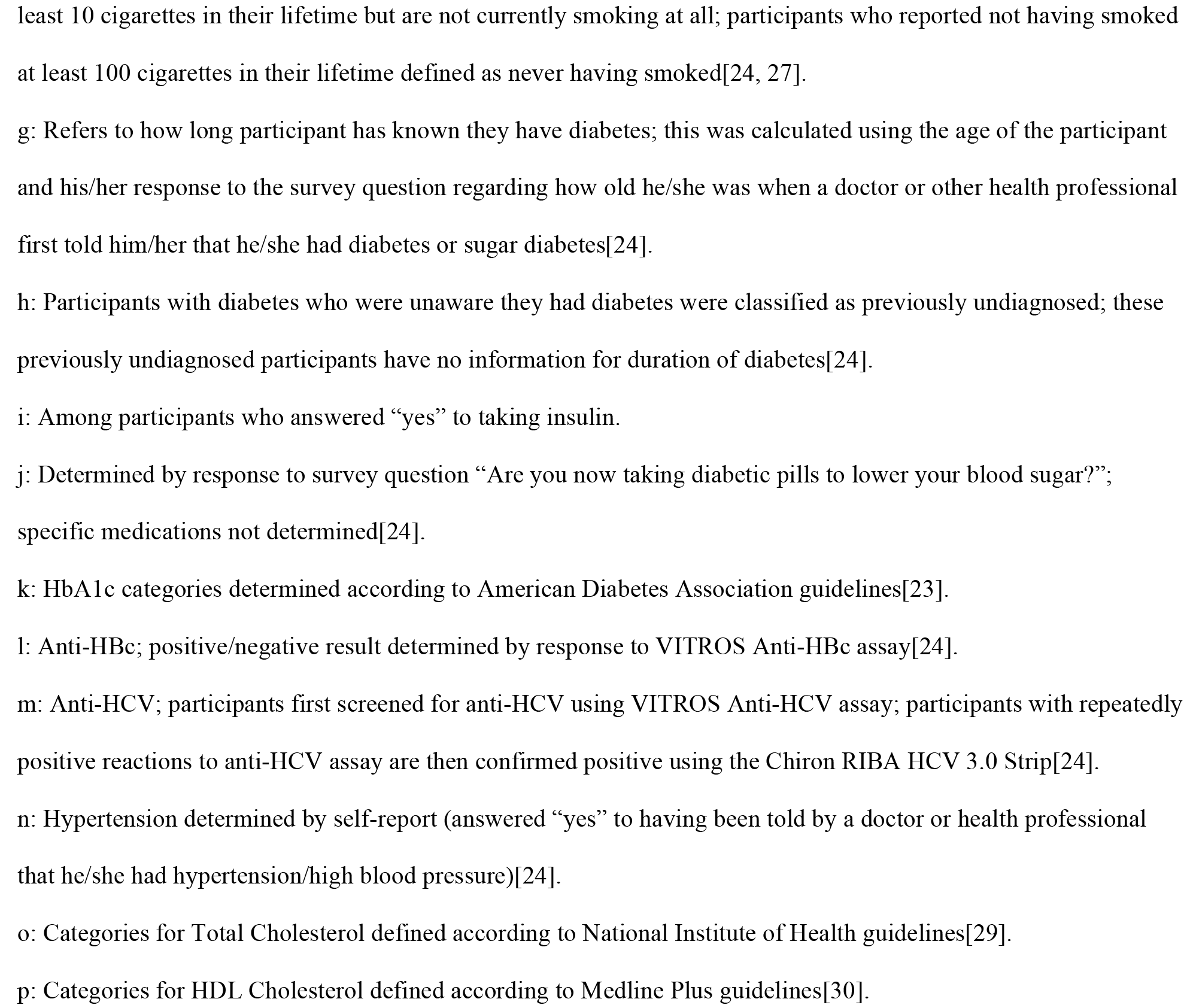
Weighted prevalence of latent tuberculosis (LTBI) infection among only those with diabetes^a^ in the civilian, non-institutionalized United States population, adults 20 years and older, 2011-2012

### 1.3.5 Sensitivity Analyses

In our sensitivity analysis to assess potential misclassification of diabetes using FBG in addition to self-report and HbA1c, adults with diabetes had significantly higher crude odds of LTBI (crude OR 2.43; 95%CI 1.32-4.49) compared to those without diabetes (data not shown). Adults with prediabetes had non-significantly higher crude odds of LTBI (crude OR 1.21; 95%CI 0.74-2.00) compared to those without diabetes. After adjusting for age, sex, smoking status, history of active TB, and foreign born status, adults with diabetes had a non-significant higher odds of LTBI (aOR 1.36; 95%CI 0.70-2.65) compared to those without diabetes.

In our sensitivity analysis to assess covariate misspecification of adjusted models, we found adjusted odds ratios that ranged from 1.49 (95%CI 0.83-2.68) to 2.20 (95%CI 1.22-3.96) for the odds of LTBI in adults with diabetes compared to those without diabetes (Supplemental Table 1). We found adjusted odds ratios that ranged from 0.95 (95%CI 0.75-1.21) to 1.25 (95%CI 0.93-1.68) for the odds of LTBI in adults with prediabetes compared to those without diabetes; however, none were statistically significant.

## 1.4 Conclusions

We used data nationally representative of the US population to examine the association between LTBI and diabetes and found a robust relationship between the two diseases. We reported that the prevalence of LTBI among adults with diabetes was more than twice the prevalence of those without diabetes. Similarly, we found that more than one-fifth of adults with LTBI had diabetes. We did not find significant differences in LTBI prevalence among those who were previously diagnosed compared to those who were previously undiagnosed. We also did not find significant differences in LTBI prevalence among those with prediabetes compared to those without diabetes. To our knowledge, this study is the largest and most generalizable analysis to compare the prevalence of LTBI among adults with and without diabetes and prediabetes.

Our results are consistent with the findings of previous studies. In a systematic review conducted by Lee et al., the meta-analysis included findings from one cohort study and 12 crosssectional studies investigating the association between diabetes and LTBI. From the 12 crosssectional studies, researchers calculated a pooled odds ratio of 1.18 (95%CI 1.06-1.30), indicating a slight yet significant increased odds of LTBI among patients with diabetes compared to patients without diabetes[17]. A limitation of several studies reviewed in Lee et al.’s systematic review and meta-analysis was the potential misclassification of LTBI due to measurement error associated with the TST (tuberculin skin test). Unlike many previous studies, our study relied upon the use of QFT-GIT which is not affected by the Bacillus Calmette-Guérin (BCG) vaccine. Our national estimates of LTBI prevalence were similar to previously reported estimates[18, 34].

Our results were similar to a study conducted by Hensel et al. in the metropolitan area of Atlanta, Georgia[5]. This study also utilized both HbA1c and QFT-GIT to determine diabetes status and LTBI status, respectively. Hensel et al. found a nearly doubled prevalence of LTBI among patients with diabetes compared to those without diabetes[5]. Unlike our study, however, the Atlanta study was not generalizable to the US adult population, as it only included recently arrived refugees to the US[5]. As with the Atlanta study, we found no significant difference in the prevalence of LTBI among patients with previously undiagnosed diabetes compared to those with previously diagnosed diabetes[5].

Although the causal mechanisms that result in increased co-occurrence of LTBI and diabetes remain to be definitively established, there are relevant biologic hypotheses that may explain how diabetes may increase the risk of LTBI and vice-versa. Some LTBI granulomas on the spectrum of high MTB activity include bacterial replication and likely result in proximal immune signaling, a phenomena which may persist in adipose tissue[35]. Secretion of pro-inflammatory adipokines and cytokines within adipocytes could interfere with insulin regulation and contribute to diabetes risk[36]. If LTBI contributes to immune activation within visceral adipose tissue, it would likely increase the risk of diabetes or prediabetes. Alternatively, chronic low-grade inflammation and immunopathology associated with diabetes and prediabetes [37] may contribute to susceptibility to TB infection[9, 10].

Our study was subject to several limitations. First, there may have been misclassification of participant characteristics. For example, self-reported information on smoking status was determined via participant responses to a questionnaire, so smokers may have reported not smoking due to social stigma. While diabetes and LTBI status may also be subject to misclassification, we defined these primary study variables using currently recommended clinical measures (HbA1c and QFT-GIT)[38, 39]. By using the QFT-GITs instead of the TST to measure LTBI, we avoided potential cross-reaction with antigens found in the BCG vaccine, commonly used outside the United States [39]. However, we did not account for discordance between QFT-GIT and TST, and therefore some misclassification of LTBI may have occurred. Second, in this study we were unable to adjust for the probability being exposed to someone with active TB. Although previous history of active TB was assessed via questionnaire and found to be associated with LTBI but not diabetes status, the inability to adjust for probability of exposure to TB may have distorted our estimated association between LTBI and diabetes. Nonetheless, we did adjust for several other key confounding factors such as smoking, age, and foreign born status. We also were able to assess the distribution of other underlying infections, such as hepatitis B and C and kidney disease, and found no evidence of confounding. Third, our study was a cross-sectional design, and as such we were unable to determine the temporal relationship between LTBI and diabetes. For example, our results are unable to differentiate whether the observed association was due to an increased risk of LTBI from diabetes or if LTBI may increase the risk of diabetes. Longitudinal studies are needed to investigate the temporal association between LTBI and diabetes.

This study reported that diabetes was significantly associated with an increased odds of LTBI prevalence in US adults, even after adjusting for key confounding factors. Overall, more than one-fifth of all adults with LTBI had diabetes. Information from this study greatly improves our understanding of the intersection of the TB and diabetes epidemics. With the increasing prevalence of diabetes in areas with the highest burden of TB, targeted efforts may be needed to address the co-infection of diabetes and LTBI to prevent an increase in TB incidence worldwide.

**Figure 1:**
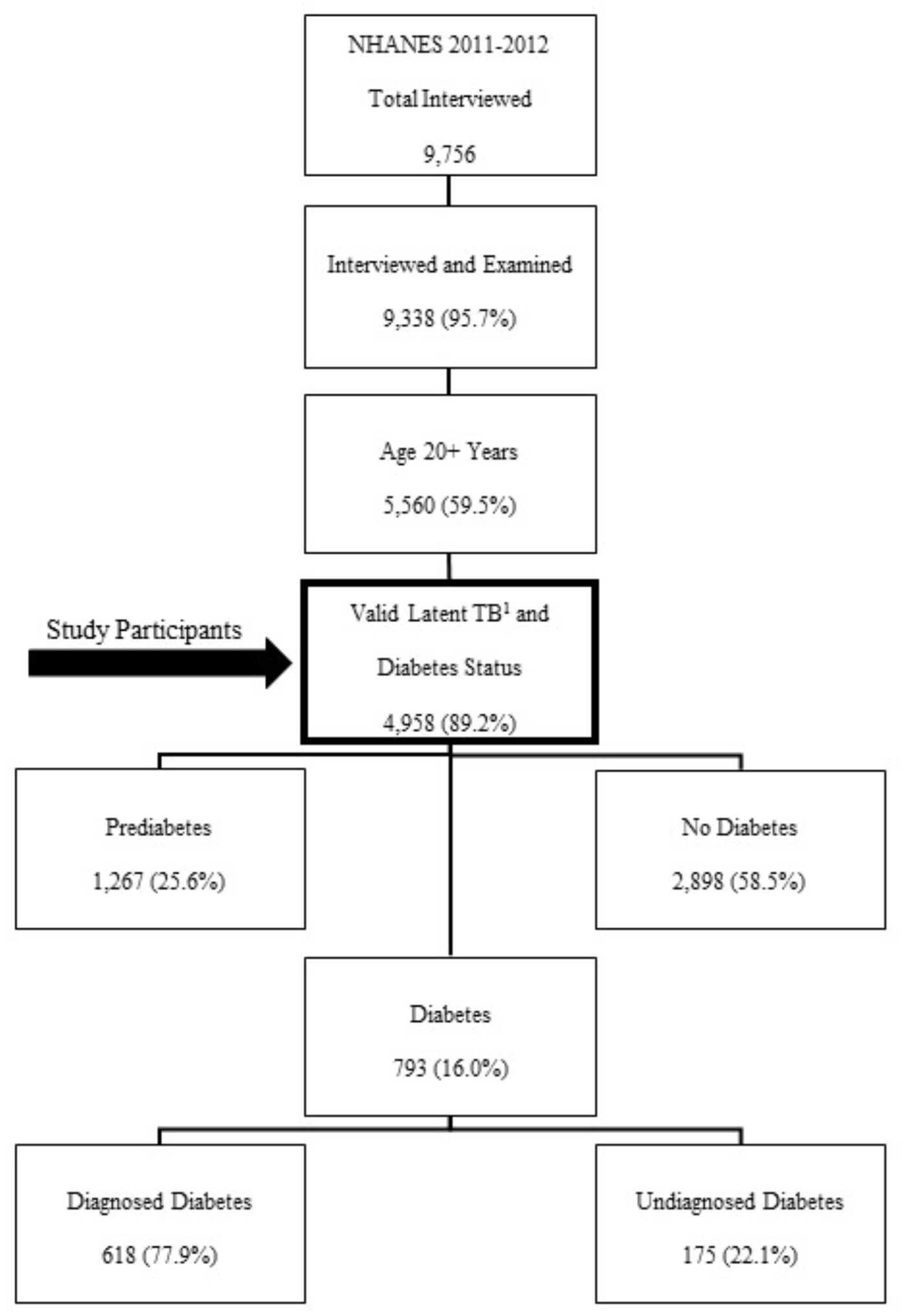
Flow chart showing process of selection for NHANES 2011-2012 participants eligible for study, including the categorization of eligible participants by diabetes status; raw numbers and percentages not weighted for NHANES sampling methodology. 1: Valid latent TB status based on QFT-GIT result

## 1.5 Acknowledgements

The findings and conclusions in this report are those of the authors and do not necessarily represent the official position of the Centers for Disease Control and Prevention (CDC). The use of trade names and commercial sources is for identification only and does not imply endorsement by the CDC. Author contributions: MJM, MMB, KMB, and MKA conceived and designed the study and drafted the initial manuscript. MMB, KMS, and MJM performed the data analyses. All authors contributed to interpretation of the data, revised the manuscript, and approved the final version.

This research did not receive any specific grant from funding agencies in the public, commercial, or not-for-profit sectors.

